# Blood flow modeling under LVAD physiology. From global circulation to local hemodynamics^☆^

**DOI:** 10.1101/2022.01.28.478161

**Authors:** Pablo J. Blanco, Jonathan Grinstein, Carlos A. Bulant, Ryo Torii, Christos V. Bourantas, Pedro A. Lemos, Héctor M. García-García

## Abstract

This document presents the modeling strategy to address the in-silico study of different LVAD patho-physiological scenarios. The proposed approach employs a closed-loop lumped-parameter compartmental representation of the global circulation in the cardiovascular system (CVS). The CVS is coupled to a HeartMate 3 LVAD, and different cardiovascular conditions are proposed by modification of model parameters. Once the simulation for these conditions are performed, the cardiac function is analyzed in detail, and the global circulation model delivers flow rate waveforms which are employed as boundary conditions in a 3D hemodynamic simulation. This local circulation model is built using a patient-specific geometry of the aortic arch, containing 7 inlet/outlet boundaries, namely: LVAD cannula, aortic root, left and right subclavian arteries, left and right common carotid arteries and thoracic aorta. This model is exploited to investigate the impact of global cardiovascular conditions in the local hemodynamic features, particularly the wall shear stress (WSS) in different spatial regions.

## 1. Introduction

The methodology is split into two phases. First, we built a closed-loop model of the cardiovascular system to analyze global circulatory phenomena, with emphasis in the cardiac performance and its interaction with the LVAD. Second, we retrieved the hemodynamic conditions, specifically blood flow rates through the LVAD cannula and aortic root, and used them as boundary conditions to perform 3D simulations using a patient-specific geometric model of the aortic arch.

The cardiovascular model developed in the present study has been specifically tailored for the present application of cardiovascular physiology coupled to LVAD function. Nevertheless, such a model shares many components with models available in the literature. Table 1 presents the supporting bibliographic works that have been employed in the instantiation of both the global and local circulation models. These bibliographic references share similar modeling ingredients, and they served to guide the parameter selection in the present investigation.

**Table 1:** Supporting bibliographic references for chosen techniques and parameter selection in the different components of the present model.

| Global circulation model |  |  |
| --- | --- | --- |
| Ingredient | Section | References |
| Closed loop representation | 2.1 | [1, 2] |
| Cardiac chamber equations and parameters | 2.2 | [1, 2, 3] |
| Valve equations and parameters | 2.3 | [1, 4, 5] |
| Arterial blood flow equations | 2.4 | [1, 2] |
| Venous blood flow equations | 2.5 | [1, 2] |
| Pulmonary blood flow equations | 2.6 | [1, 2] |
| Post-processing | 2.9 | [6] |
| Local circulation model |  |  |
| 3D Model geometry | 3.1 | [7] |
| 3D Model equations | 3.2 | [8, 9] |
| Post-processing | 3.4 | [8] |

## 2. Global circulation model

### 2.1. Closed-loop representation and notation

The global circulation model consists of a set of inter-connected compartments whose dynamics is characterized by a set of ordinary differential equations. Figure 1 schematically illustrates the compartments that form the closed-loop representation of the CVS.

**Figure 1:**
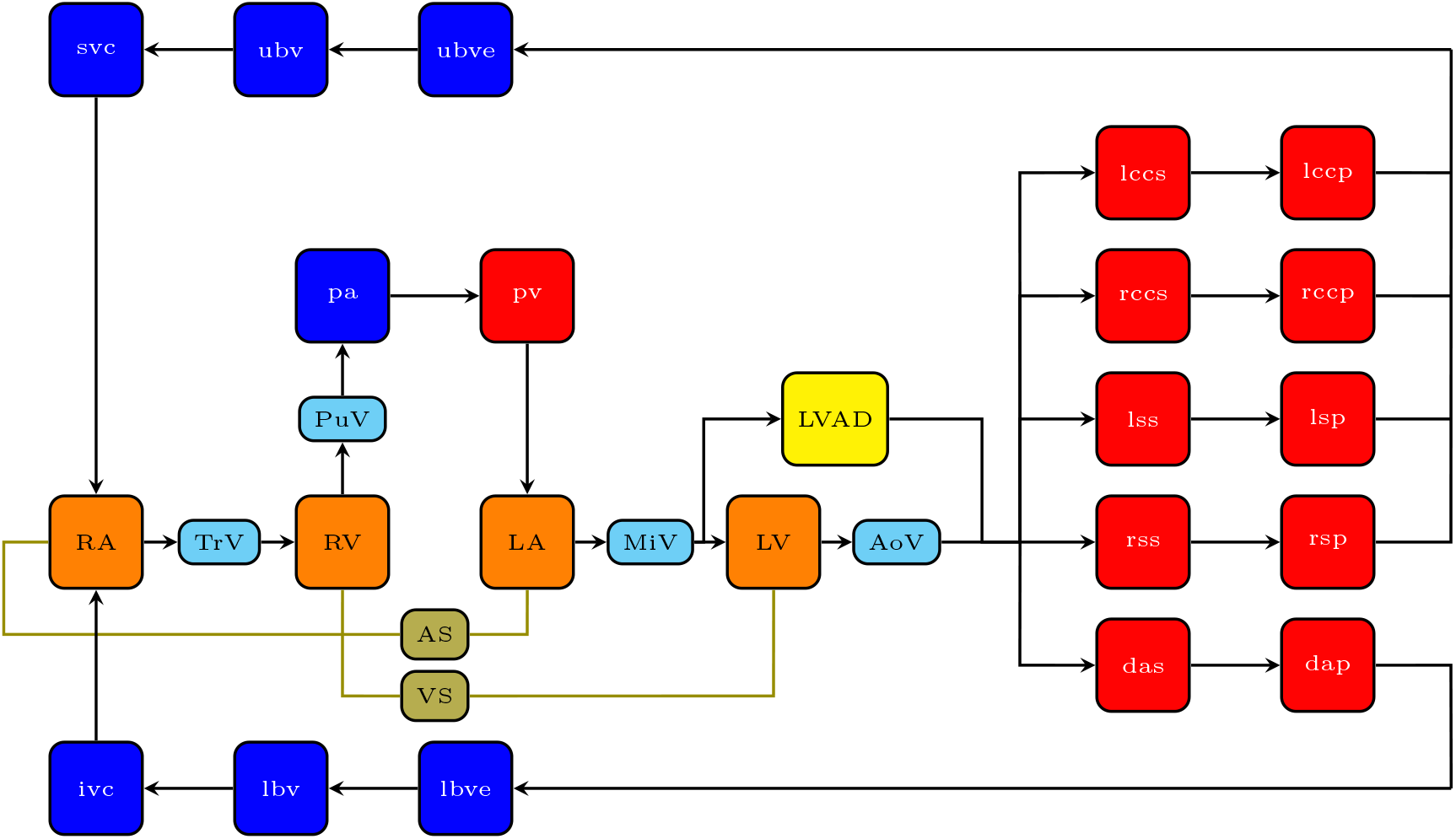
Closed-loop model of the CVS. Red: arterial compartments. Blue: venous compartments. Light blue: valve compartmens. Orange: cardiac compartments. Lime: septal compartments. Yellow: LVAD compartment.

The main common variables across the closed-loop system are the blood pressure, denoted by *P*, and the flow rate, denoted by *Q*. In the cardiac chambers, and in the cardiac valves, chamber volume, denoted by *V*, and valvel opening angle, denoted by *θ*, are also described by the system of equations

In the system illustrated in Figure 1, the following notational convention is used concerning indexes

- RA, LA, RV, LV: right/left atria, right/left ventricles
- AS, VS: atrial septum, ventricular septum
- TrV, PuV, MiV, AoV: tricuspid, pulmonary, mitral, aortic valves
- pa, pv: pulmonary arteries/veins
- LVAD: left ventricular assist device
- lccsi, rccsi, lccs, rccs, lccp, rccp: left/right common carotid systemic inlet, left/right common carotid systemic, left/right common carotid peripheral
- lssi, rssi, lss, rss, lsp, rsp: left/right subclavian systemic inlet, left/right subclavian systemic, left/right subclavian peripheral
- dasi, das, dap: descending aorta systemic inlet, descending aorta systemic, descending aorta peripheral
- ubve, lbve, ubv, lbv, svc, ivc: upper/lower body venules, upper/lower body veins, superior/inferior vena cava
- u, d: upstream and downstream (circulation-wise), respectively; these indexes are used to characterize connectivity with upstream and downstream components

The model descriptive capabilities in terms of hemodynamic variables are described in Table 2, including pressure, flow rate quantities, chamber volume and valve opening angle unknowns. Table 3 describes the connectivities across the entire closed-loop system, which allow to write the equations to be presented below in a more compact way. In turn, the list of parameters, including their specific physical significance, is presented in Section 4.

**Table 2:**
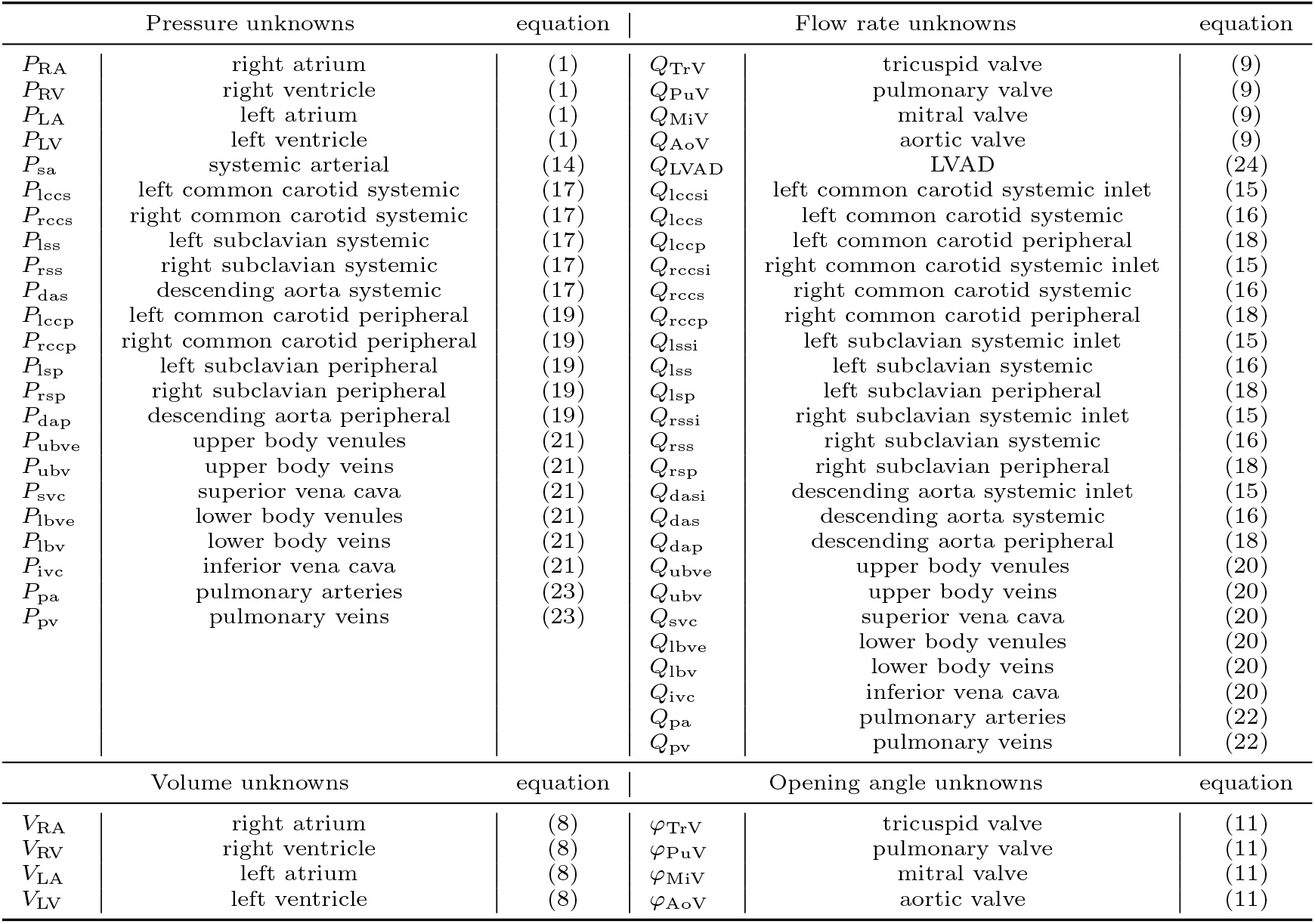
Unknowns corresponding to the blood pressure, flow rate, chamber volume and valve opening angle across the different compartments of the closed-loop CVS system.

**Table 3:** Upstream and downstream connectivities for pressure and flow rate variables in the closed-loop system from Figure 1.

| Quantity | connectivity x |  |
| --- | --- | --- |
|  | u | d |
| $P_{TrV}^x$ | $P_{RA}$ | $P_{RV}$ |
| $P_{PuV}^x$ | $P_{RV}$ | $P_{pa}$ |
| $P_{MiV}^x$ | $P_{LA}$ | $P_{LV}$ |
| $P_{AoV}^x$ | $P_{LV}$ | $P_{sa}$ |
| $P_{lccs}^x$ | $P_{lccs}$ | $P_{lccp}$ |
| $P_{rccs}^x$ | $P_{rccs}$ | $P_{rccp}$ |
| $P_{lss}^x$ | $P_{lss}$ | $P_{lsp}$ |
| $P_{rss}^x$ | $P_{rss}$ | $P_{rsp}$ |
| $P_{das}^x$ | $P_{das}$ | $P_{dap}$ |
| $P_{lccp}^x$ | $P_{lccp}$ | $P_{ubve}$ |
| $P_{rccp}^x$ | $P_{rccp}$ | $P_{ubve}$ |
| $P_{lsp}^x$ | $P_{lsp}$ | $P_{ubve}$ |
| $P_{rsp}^x$ | $P_{rsp}$ | $P_{ubve}$ |
| $P_{dap}^x$ | $P_{dap}$ | $P_{lbve}$ |
| $P_{ubve}^x$ | $P_{ubve}$ | $P_{ubv}$ |
| $P_{ubv}^x$ | $P_{ubv}$ | $P_{svc}$ |
| $P_{svc}^x$ | $P_{svc}$ | $P_{RA}$ |
| $P_{lbve}^x$ | $P_{lbve}$ | $P_{lbv}$ |
| $P_{lbv}^x$ | $P_{lbv}$ | $P_{ivc}$ |
| $P_{ivc}^x$ | $P_{ivc}$ | $P_{RA}$ |
| $P_{pa}^x$ | $P_{pa}$ | $P_{pv}$ |
| $P_{pv}^x$ | $P_{pv}$ | $P_{LA}$ |
|  | u | d |
| $Q_{RA}^x$ | $Q_{svc} + Q_{ivc}$ | $Q_{TrV}$ |
| $Q_{RV}^x$ | $Q_{TrV}$ | $Q_{PuV}$ |
| $Q_{LA}^x$ | $Q_{pv}$ | $Q_{MiV}$ |
| $Q_{LV}^x$ | $Q_{MiV}$ | $Q_{AoV}$ |
| $Q_{lccs}^x$ | $Q_{lccsi}$ | $Q_{lccs}$ |
| $Q_{rccs}^x$ | $Q_{rccsi}$ | $Q_{rccs}$ |
| $Q_{lss}^x$ | $Q_{lssi}$ | $Q_{lss}$ |
| $Q_{rss}^x$ | $Q_{rssi}$ | $Q_{rss}$ |
| $Q_{das}^x$ | $Q_{dasi}$ | $Q_{das}$ |
| $Q_{lccp}^x$ | $Q_{lccs}$ | $Q_{lccp}$ |
| $Q_{rccp}^x$ | $Q_{rccs}$ | $Q_{rccp}$ |
| $Q_{lsp}^x$ | $Q_{lss}$ | $Q_{lsp}$ |
| $Q_{rsp}^x$ | $Q_{rss}$ | $Q_{rsp}$ |
| $Q_{dap}^x$ | $Q_{das}$ | $Q_{dap}$ |
| $Q_{ubve}^x$ | $Q_{lccp} + Q_{rccp} + Q_{lsp} + Q_{rsp}$ | $Q_{ubve}$ |
| $Q_{ubv}^x$ | $Q_{ubve}$ | $Q_{ubv}$ |
| $Q_{svc}^x$ | $Q_{ubv}$ | $Q_{svc}$ |
| $Q_{lbve}^x$ | $Q_{dap}$ | $Q_{lbve}$ |
| $Q_{lbv}^x$ | $Q_{lbve}$ | $Q_{lbv}$ |
| $Q_{ivc}^x$ | $Q_{lbv}$ | $Q_{ivc}$ |
| $Q_{pa}^x$ | $Q_{PuV}$ | $Q_{pa}$ |
| $Q_{pv}^x$ | $Q_{pa}$ | $Q_{pv}$ |

### 2.2. Cardiac chamber equations

The pressure-volume relation for the cardiac chambers are the following

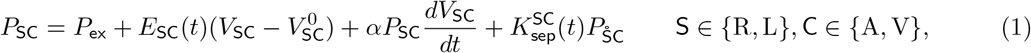

where 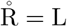 and 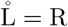.

The cardiac elastances are defined by means of proper activation functions, and are coupled through the septal interaction

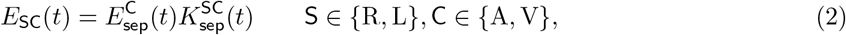

where the septal inter-atria and inter-ventricular interactions are defined as follows

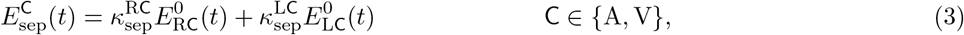

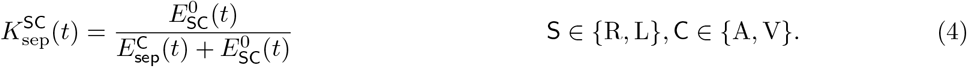

The native chamber elastances are given by

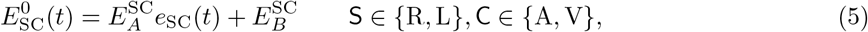

where the atrial and ventricular activation functions are

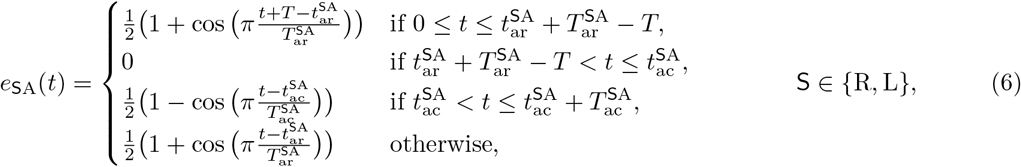

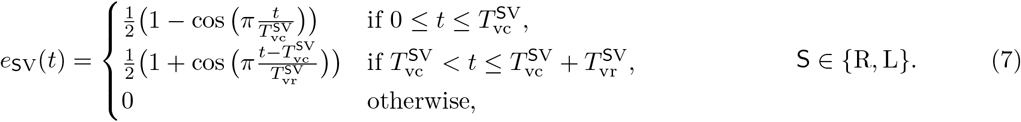

The mass conservation equations in the cardiac chambers read

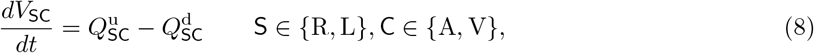

where the upstream and downstream connectivities are given in Table 3.

### 2.3. Valve equations

The governing equations for the blood flow across the cardiac valves are the following (note the constraints defined in (13)

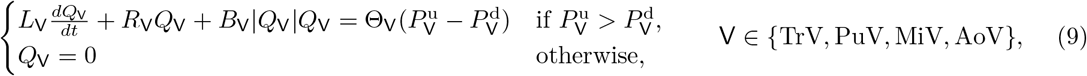

where the orifice coefficients are defined through the opening angles as follows

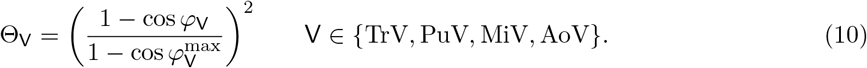

In turn, the governing equations for the valve opening angles are the following

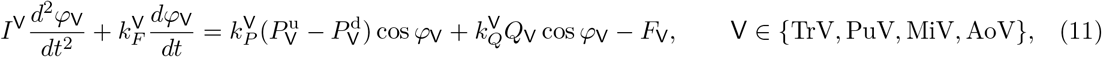

where

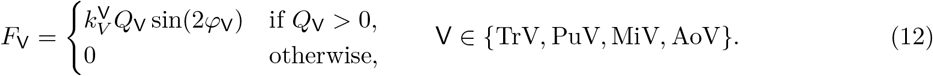

Finally, the valve opening angle is constrained to be in a certain range, as follows

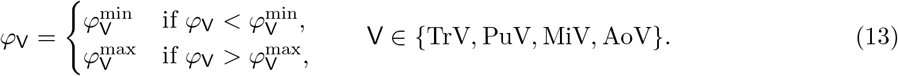

### 2.4. Arterial blood flow equations

At the entrance of the arterial system, where the LVAD and the aortic root discharge blood into the five systemic territories, the following equations hold

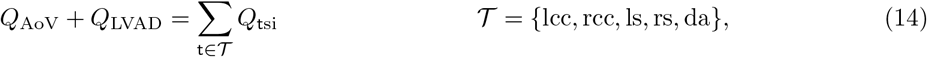

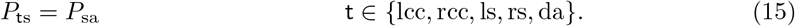

Conservation of momentum in systemic arterial compartments is governed by the following equations

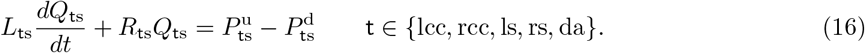

Mass conservation in the systemic territories is governed by the following equations

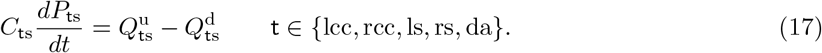

The five peripheral beds, connected to the corresponding systemic compartmens, are modeled using similar expressions. Thas is, for the conservation of momentum

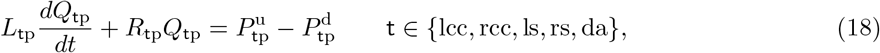

and for the conservation of mass

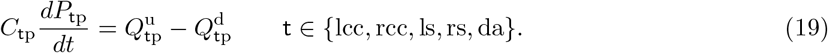

### 2.5. Venous blood flow equations

The blood flow equations for the venous compartments are analogous to those employed in the arterial side. For the upper and lower body regions, the following momentum conservation equations for each of the six compartments are employed

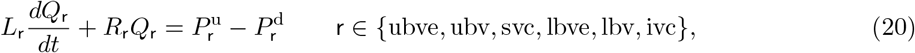

and for the mass conservation it is

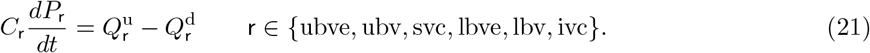

### 2.6. Pulmonary blood flow equations

Conservation of momentum in the pulmonary arteries and veins is governed by the following equations

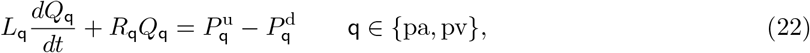

and the corresponding equations for the mass conservation are

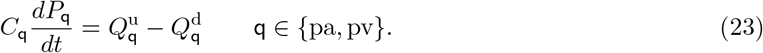

### 2.7. LVAD performance

The LVAD performance is characterized through a pressure-flow rate relation corresponding to a HeartMate 3 device. This non-linear function is expressed as follows

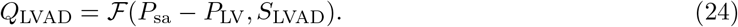

Starting from a set of pairs 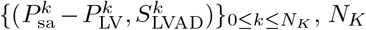 the number of data points, a regression strategy was utilized to model the relation ℱ, and the resulting relations are shown in Figure 2.

**Figure 2:**
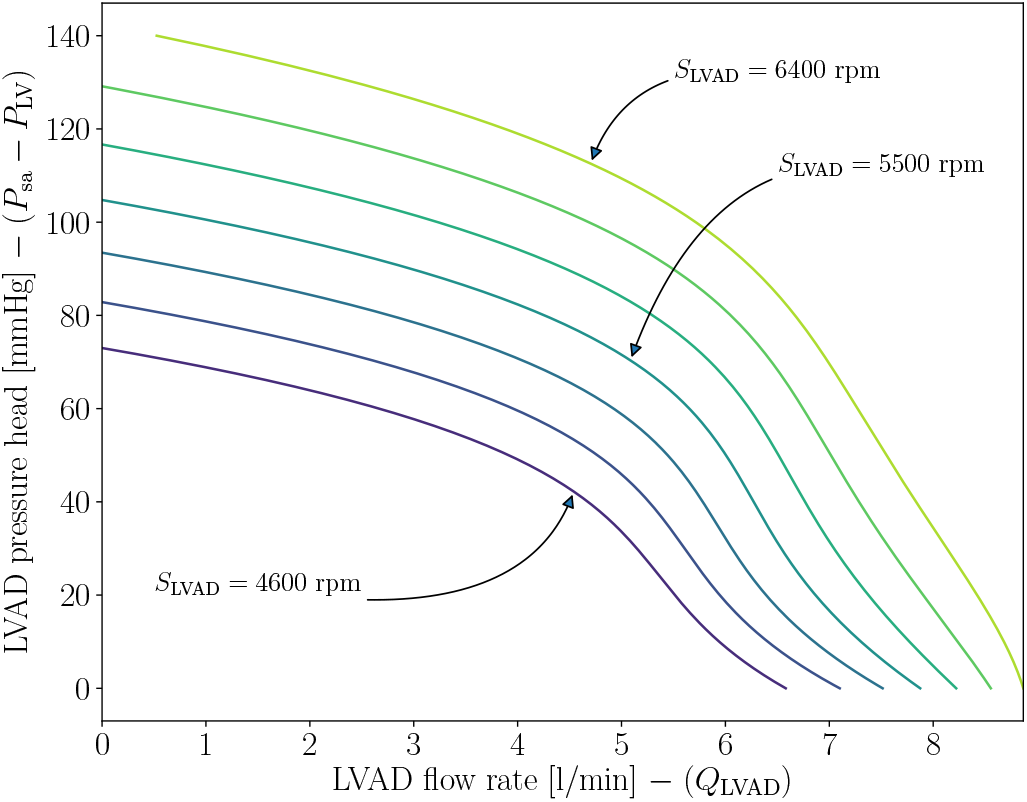
Pressure-flow rate relations at different LVAD speeds. These curved correspond to a HeartMate 3 device. There is a jump of 300 rpm in LVAD speed between consecutive curves.

### 2.8. Simulation

The initial condition for the model is a physiological initial guess, and a total of ten cardiac cycles is run to guarantee that the model reached a periodic regime.

The global circulation model is discretized in time using a fully implicit backward Euler scheme. Non-linearities are treated using Picard iterations. The discrete model describes the temporal dynamimcs of the 59 unknowns in the problem. At each time-step, the resulting linear system of algebraic equations is solved using LU factorization. The global circulation model is run in a laptop, and takes less than 2 minutes to simulate 10 cardiac cycles.

### 2.9. Post-processing

Once the simulations are finished, we take the last cardiac cycle and compute several quantities of interest that help to characterize the cardiac function and the cardiovascular system as a whole. The list of quantities of interest is given in Table 4. Figure 3 illustrates some PV loops for RV and LV, with the corresponding landmarks that play a role in the processing of cardiac variables.

**Table 4:** List of properties that characterize the cardiac function and the cardiovascular system.

| left/right ventricle (SV, S ∈ {R, L}) - computed using P<sub>SV</sub>-V<sub>SV</sub> loop |  |  |
| --- | --- | --- |
| description | symbol | equation |
| ventricular end-systolic elastance | $E_{esSV}$ | (25) |
| effective pulmonary/systemic arterial elastance | $E_{aSV}$ | (28) |
| ventricular-arterial elastance ratio | $F_{SV}^{es/a}$ | (29) |
| ventricular stroke work | $SW_{SV}$ | (30) |
| end-systolic potential energy | $PE_{SV}$ | (31) |
| pressure-volume area | $PVA_{SV}$ | (32) |
| myocardial O<sub>2</sub> consumption | $MVO_{2SV}$ | (33) |
| ventricular efficiency | $Eff_{SV}$ | (34) |

| pressure and flow rate indicators<br>obtained from querying model variables |  |  |
| --- | --- | --- |
| description | symbol | equation |
| mean systemic arterial pressure | $\bar{P}_{sa}$ | (35) |
| systolic systemic arterial pressure | $P_{sa}^{max}$ | (36) |
| diastolic systemic arterial pressure | $P_{sa}^{min}$ | (37) |
| mean pulmonary arterial pressure | $\bar{P}_{pa}$ | (38) |
| systolic pulmonary arterial pressure | $P_{pa}^{max}$ | (39) |
| diastolic pulmonary arterial pressure | $P_{pa}^{min}$ | (40) |
| mean transvalvular pressure gradient | $\Delta P_{AoV}$ | (41) |
| mean aortic flow rate | $\bar{Q}_{AoV}$ | (42) |
| mean LVAD flow rate | $\bar{Q}_{LVAD}$ | (43) |
| volumetric aortic backflow | $V_{AoV}^{back}$ | (45) |
| regurgitant fraction | $RF$ | (46) |

**Figure 3:**
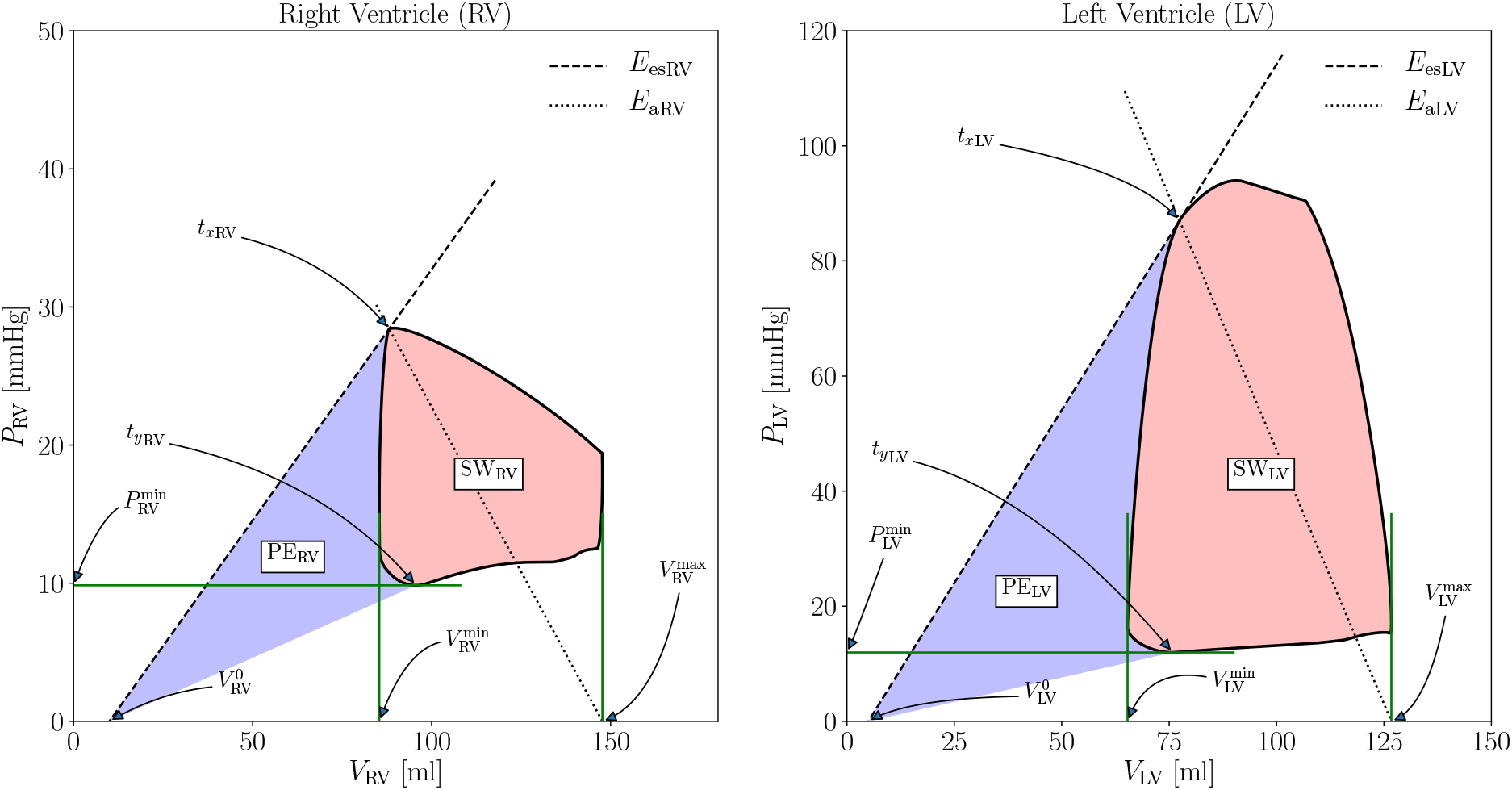
Prototypical pressure-volume loops for RV and LV, with the corresponding landmark points that play a role in the characterization of the cardiac function.

From the RV and LV pressure-volume loops we compute the following quantities for S ∈ {R, L}

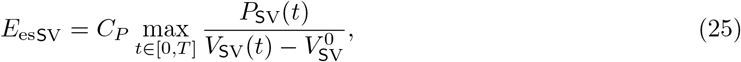

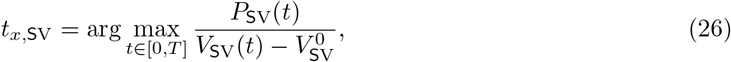

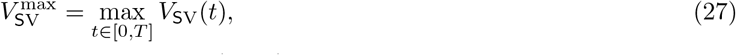

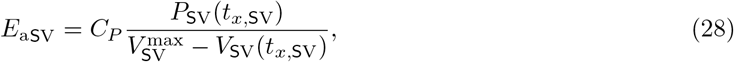

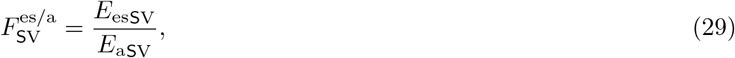

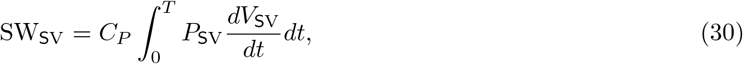

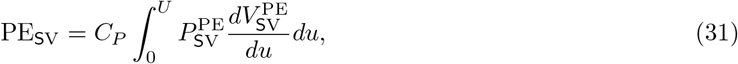

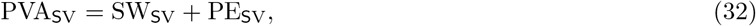

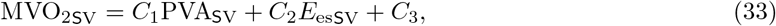

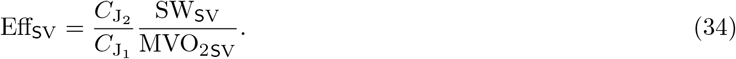

In (31), we define the parametric curves 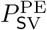 and 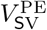 as those curves formed by connecting the points 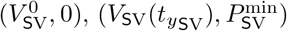 and 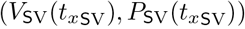 in Figure 3.

Parameters *C*_1_ and *C*_2_ in equation (33) are defined in Table 5. Also in that table, we report the parameters that play a role in the unit conversion process.

**Table 5:** Parameters that characterize the computation of derived quantities of interest in the analysis of the cardiac function.

| description | symbol | value | units |
| --- | --- | --- | --- |
| pressure unit transformation coefficient | $C_P$ | 1.0/1333.32 | (mmHg)/(dyn/cm<sup>2</sup>) |
| flow rate unit transformation coefficient | $C_Q$ | 60/1000 | (l/min)/(cm<sup>3</sup>/s) |
| oxygen consumption constant | $C_1$ | $2.1 \cdot 10^{-5}$ | mlO<sub>2</sub>/(mmHg·ml) |
| oxygen consumption constant | $C_2$ | $3.2 \cdot 10^{-3}$ | (mlO<sub>2</sub>/beat) · (ml/mmHg) |
| oxygen consumption constant | $C_3$ | $1.04 \cdot 10^{-2}$ | mlO<sub>2</sub>/beat |
| Joule transformation coefficient 1 | $C_{J_1}$ | 20.0 | J/(mlO<sub>2</sub>/beat) |
| Joule transformation coefficient 2 | $C_{J_2}$ | $1.33 \cdot 10^{-4}$ | J/(mmHg·ml) |

Also, from the results, we obtain the following values, converted to units of mmHg, corresponding to the mean, maximum and minimum systemic and pulmonary arterial pressures

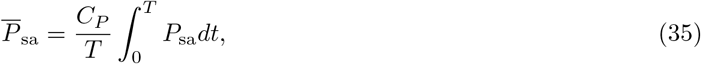

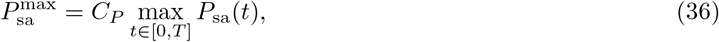

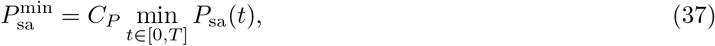

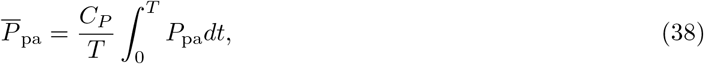

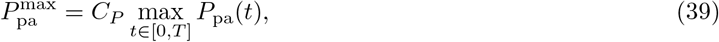

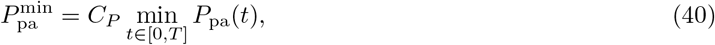

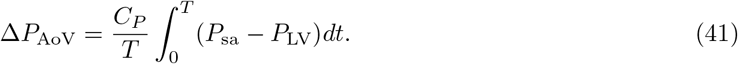

Analogously, we compute the mean flow rate through the aortic root, and through the LVAD, converted to units of l/min

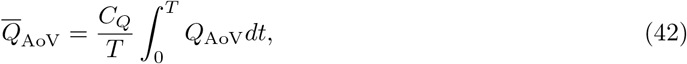

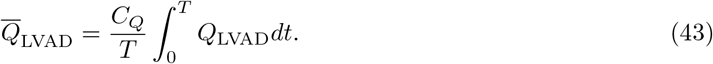

The resulting volumetric backflow through the aortic valve is computed as follows

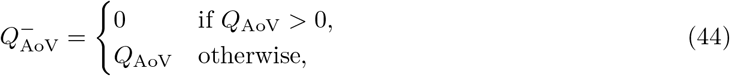

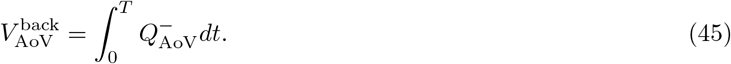

The regurgitant fraction (RF) is defined as follows

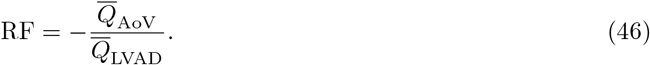

Note that since RF is defined whenever 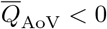, and in that case RF > 0.

## 3. Local circulation model

### 3.1. 3D Model geometry

The mathematical model to simulate the the local blood flow occurring in the proximal aorta and its branches is given by the Navier-Stokes equations. Let Ω be the vascular domain (proximal aorta including major branches) and let Γ be its boundary, which is divided into different parts as follows

- Γ_*L*_: the lateral wall (endothelial wall),
- Γ_LVAD_: the boundary corresponding to the LVAD cannula
- Γ_AoV_: the boundary corresponding to the aortic root, right after the aortic valve
- Γ_lccsi_: the boundary corresponding to the inlet of the left common carotid systemic territory
- Γ_rccsi_: the boundary corresponding to the inlet of the left common carotid systemic territory
- Γ_lssi_: the boundary corresponding to the inlet of the left subclavian systemic territory
- Γ_rssi_: the boundary corresponding to the inlet of the right subclavian systemic territory
- Γ_dasi_: the boundary corresponding to the inlet of the descending aorta systemic territory

The outward unit normal vector to the boundary is denoted by **n**. Figure 4 illustrates, over the left, the geometry of the proximal aorta and the basic local circulation model ingredients, and over the right the division of the endothelial surface into four major regions for analysis.

**Figure 4:**
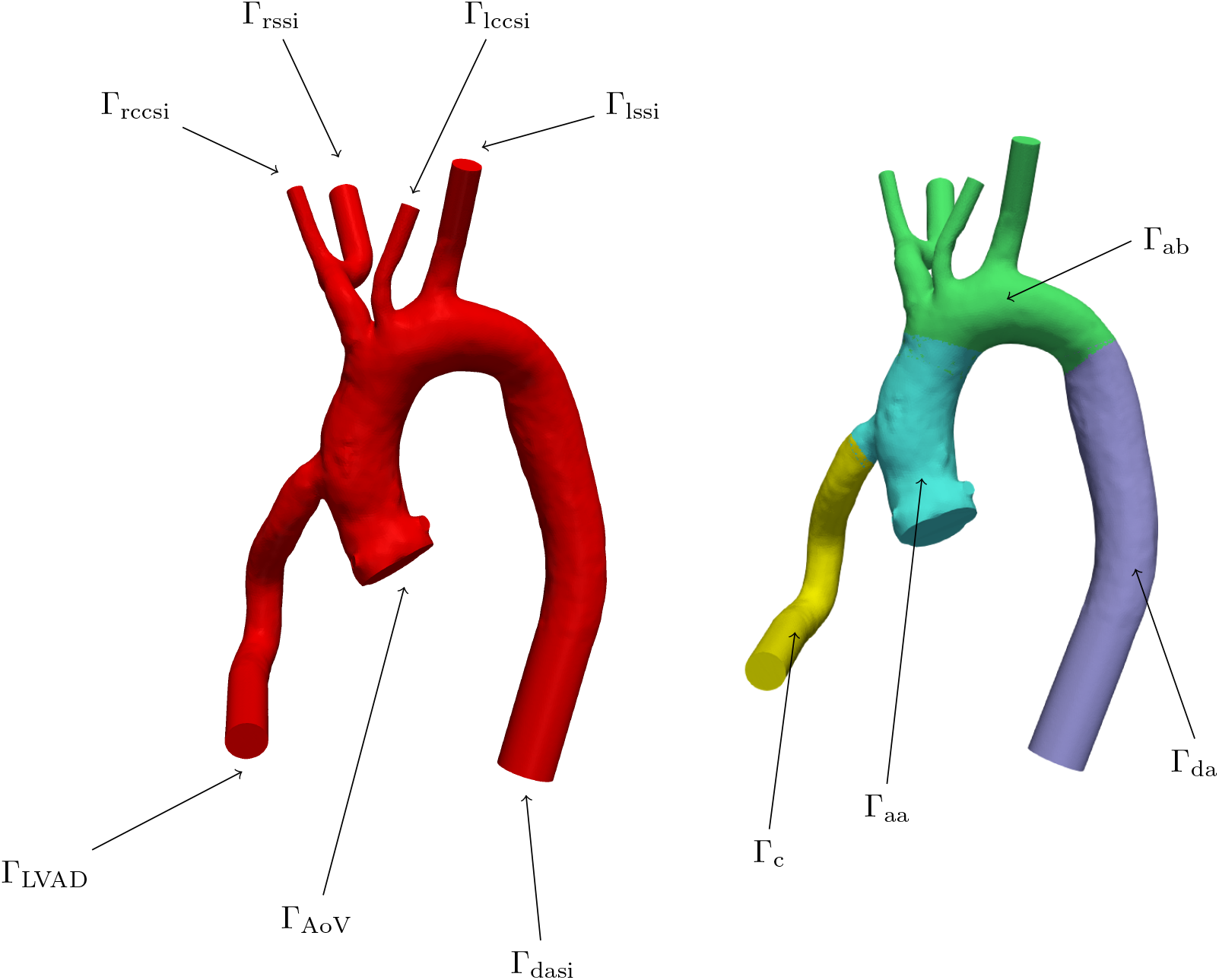
Patient-specific vascular geometry of the proximal aorta, connected to the LVAD through a 14mm cannula, with its major branches, and division of lateral endothelial wall into four major regions (c: cannula, aa: ascending aorta, ab: aortic branches and da: desdencing aorta) for analysis.

### 3.2. 3D Model equations

Let *p*(*t*, **x**) (scalar-valued field) and **v**(*t*, **x**) (vector-valued field) be the 3D dimensional functions that describe, respectively, the blood pressure and the blood velocity for all coordinate points **x** in Ω along time *t*. The blood flow is assumed to be incompressible, blood is assumed to behave as a Newtonian fluid, and we neglect the gravity and turbulence effects. The Navier-Stokes equations read

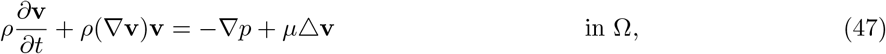

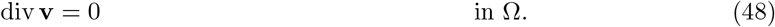

The problem is fully described once proper initial conditions and boundary conditions are prescribed. As initial conditions we prescribe null velocity

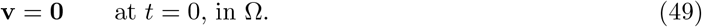

Boundary conditions must be prescribed for all the parts that describe the boundary. Over the lateral wall we assume no-slip flow. At the cannula boundary and at the aortic root, we prescribe the flow rate given by the global circulation model under the different simulation protocols that characterize the cardiovascular conditions of interest in the present study (see Section 4).

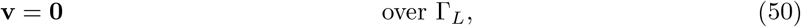

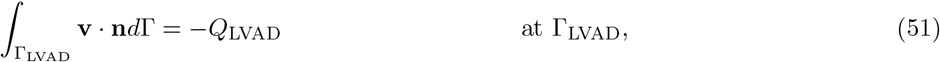

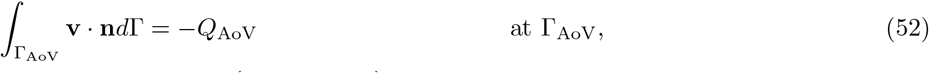

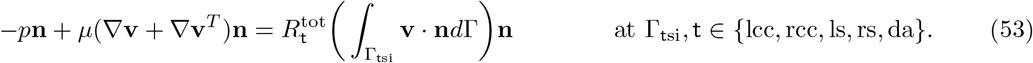

The second and third boundary conditions are called defective boundary conditions. Together with such boundary data, we assume that the traction vectors associated to the flow rate constraints are uniform at Γ_LVAD_ and at Γ_AoV_, correspondingly.

### 3.3. Simulation

The initial condition for the model is a resting condition (null blood velocity). Three cardiac cycles are simulated to ensure that the model reaches a periodic regime. The local circulation model is discretized in time using a fully implicit backward Euler scheme, and in space using the finite element method, with equal order interpolation for pressure and velocity, with the velocity space enriched through bubble functions (mini-element). Non-linearities are treated using Picard iterations. Spatial discretization is accomplished by using a tetrahedral mesh with boundary layer elements (X layers). The number of tetrahedra, and degrees of freedom in the problem are 10430016 and 6933796, respectively. The resulting algebraic system of equations is solved using GMRES method, with an overlapping additive Schwarz preconditioner. The local circulation model is run using a distributed computing paradigm in a supercomputer, using 480 computing cores, and takes on average 8 hours to simulate 1 cardiac cycle.

### 3.4. Post-processing

Once the simulations are finished, we take the velocity field **v** in the last cardiac cycle and compute the shear stress over the endothelial wall, that is

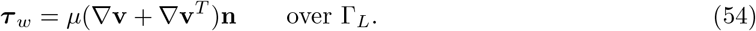

Then, we average the shear stress along the cardiac period *T*, as follows

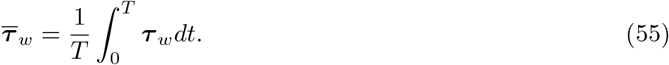

This vector field is analyzed in four large regions within the geometric model, as illustrated in Figure 4.

These regions are

- Γ_c_: cannula
- Γ_aa_: ascending aorta
- Γ_ab_: aortic branches
- Γ_da_: descending aorta

Thus, we compute the average of the time-averaged shear stress field,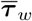, in these four endothelial surfaces, and then we compute the magnitude of the resulting vector value, that is

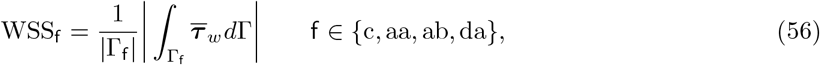

where |Γ_f_ |, denotes the endothelial area corresponding to the f region (cannula, ascending aorta, aortic branches and descending aorta).

## 4. Simulation protocols

### 4.1. Reference parameters

The entire list of reference parameters is split into several tables. These parameters are properly modified to create the different cardiovascular conditions of interest for the present study.

These are: Table 6 for cardiac chambers, Table 7 for cardiac valves, Table 8 for opening angle dynamics, Table 9 for systemic arterial territories, Table 10 for venous systemic regions, Table 11 for pulmonary arterial and venous circulations, and Table 12 for the LVAD.

**Table 6:**
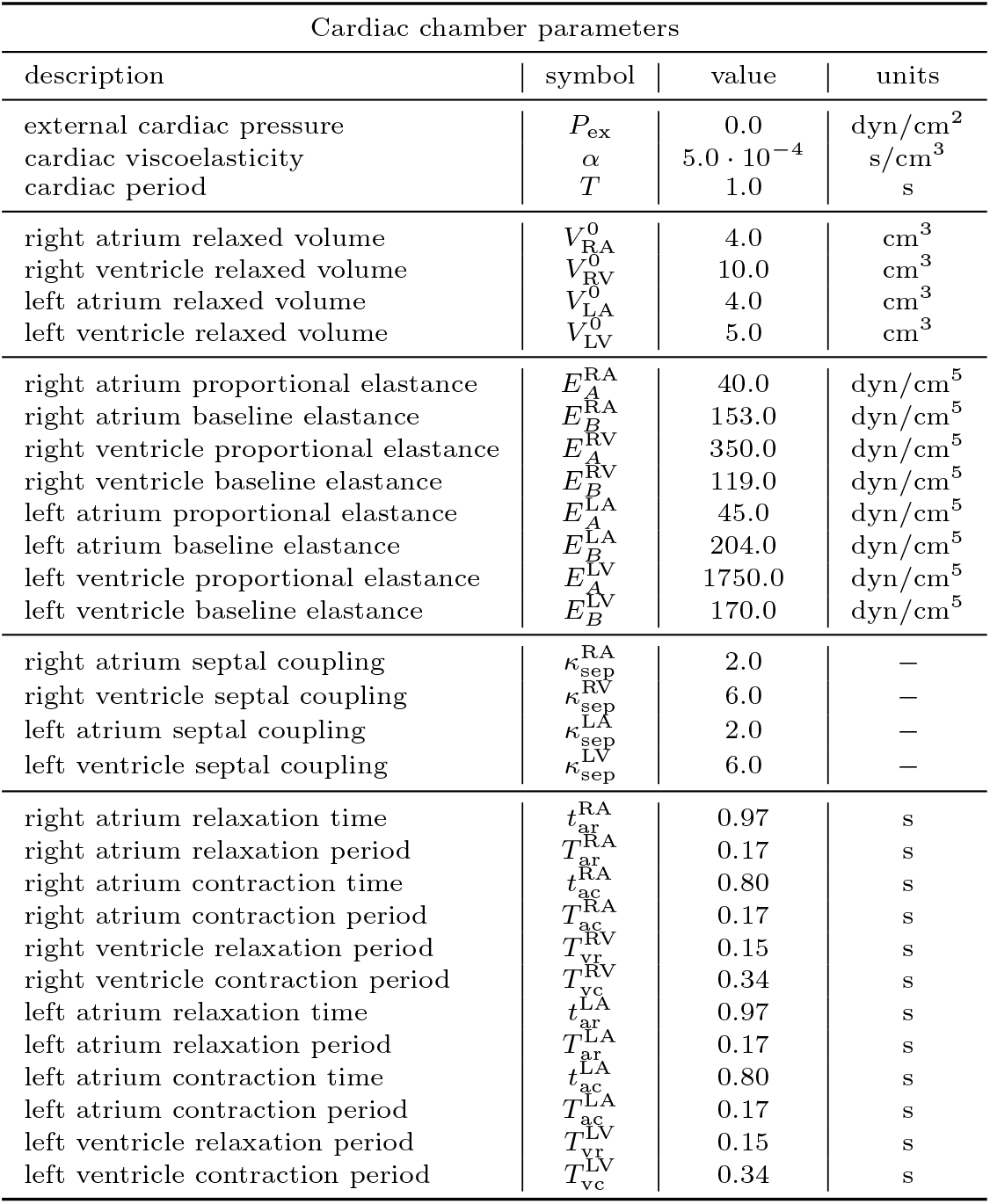
Description of model parameters that characterize the cardiac chamber compartments.

**Table 7:** Description of model parameters that characterize the cardiac valve compartments.

| Cardiac valve parameters |  |  |  |
| --- | --- | --- | --- |
| description | symbol | value | units |
| tricuspid flow inertance | $L_{TrV}$ | $5.0 \cdot 10^{-3}$ | $\text{dyn}\cdot\text{s}^2/\text{cm}^5$ |
| pulmonary flow inertance | $L_{PuV}$ | $5.0 \cdot 10^{-3}$ | $\text{dyn}\cdot\text{s}^2/\text{cm}^5$ |
| mitral flow inertance | $L_{MiV}$ | $5.0 \cdot 10^{-3}$ | $\text{dyn}\cdot\text{s}^2/\text{cm}^5$ |
| aortic flow inertance | $L_{AoV}$ | $5.0 \cdot 10^{-3}$ | $\text{dyn}\cdot\text{s}^2/\text{cm}^5$ |
| tricuspid flow resistance | $R_{TrV}$ | $6.0 \cdot 10^{-3}$ | $\text{dyn}\cdot\text{s}/\text{cm}^5$ |
| pulmonary flow resistance | $R_{PuV}$ | $6.0 \cdot 10^{-3}$ | $\text{dyn}\cdot\text{s}/\text{cm}^5$ |
| mitral flow convectance | $B_{MiV}$ | $6.0 \cdot 10^{-3}$ | $\text{dyn}\cdot\text{s}/\text{cm}^5$ |
| aortic flow resistance | $R_{AoV}$ | $6.0 \cdot 10^{-3}$ | $\text{dyn}\cdot\text{s}/\text{cm}^5$ |
| tricuspid flow convectance | $B_{TrV}$ | $6.40 \cdot 10^{-3}$ | $\text{dyn}\cdot\text{s}^2/\text{cm}^8$ |
| pulmonary flow convectance | $B_{PuV}$ | $7.56 \cdot 10^{-3}$ | $\text{dyn}\cdot\text{s}^2/\text{cm}^8$ |
| mitral flow resistance | $R_{MiV}$ | $6.40 \cdot 10^{-3}$ | $\text{dyn}\cdot\text{s}^2/\text{cm}^8$ |
| aortic flow convectance | $B_{AoV}$ | $7.56 \cdot 10^{-3}$ | $\text{dyn}\cdot\text{s}^2/\text{cm}^8$ |

**Table 8:** Description of model parameters that characterize the opening angle model.

| Opening angle parameters |  |  |  |
| --- | --- | --- | --- |
| description | symbol | value | units |
| tricuspid valve moment of inertia | $I^{TrV}$ | 0.0 | $\text{g}\cdot\text{cm}^2$ |
| pulmonary valve moment of inertia | $I^{PuV}$ | 0.0 | $\text{g}\cdot\text{cm}^2$ |
| mitral valve moment of inertia | $I^{MiV}$ | 0.0 | $\text{g}\cdot\text{cm}^2$ |
| aortic valve moment of inertia | $I^{AoV}$ | 0.0 | $\text{g}\cdot\text{cm}^2$ |
| tricuspid orifice viscosity | $k_F^{TrV}$ | 5.0 | $\text{g}\cdot\text{cm}^2/\text{s}$ |
| pulmonary orifice viscosity | $k_F^{PuV}$ | 5.0 | $\text{g}\cdot\text{cm}^2/\text{s}$ |
| mitral orifice viscosity | $k_F^{MiV}$ | 5.0 | $\text{g}\cdot\text{cm}^2/\text{s}$ |
| aortic orifice viscosity | $k_F^{AoV}$ | 5.0 | $\text{g}\cdot\text{cm}^2/\text{s}$ |
| tricuspid orifice pressure | $k_P^{TrV}$ | 50.0 | $\text{cm}^3$ |
| pulmonary orifice pressure | $k_P^{PuV}$ | 50.0 | $\text{cm}^3$ |
| mitral orifice pressure | $k_P^{MiV}$ | 50.0 | $\text{cm}^3$ |
| aortic orifice pressure | $k_P^{AoV}$ | 50.0 | $\text{cm}^3$ |
| tricuspid orifice acceleration 1 | $k_Q^{TrV}$ | 2.0 | $\text{g}/\text{cm}/\text{s}$ |
| pulmonary orifice acceleration 1 | $k_Q^{PuV}$ | 2.0 | $\text{g}/\text{cm}/\text{s}$ |
| mitral orifice acceleration 1 | $k_Q^{MiV}$ | 2.0 | $\text{g}/\text{cm}/\text{s}$ |
| aortic orifice acceleration 1 | $k_Q^{AoV}$ | 2.0 | $\text{g}/\text{cm}/\text{s}$ |
| tricuspid orifice acceleration 2 | $k_V^{TrV}$ | 3.5 | $\text{g}/\text{cm}/\text{s}$ |
| pulmonary orifice acceleration 2 | $k_V^{PuV}$ | 3.5 | $\text{g}/\text{cm}/\text{s}$ |
| mitral orifice acceleration 2 | $k_V^{MiV}$ | 3.5 | $\text{g}/\text{cm}/\text{s}$ |
| aortic orifice acceleration 2 | $k_V^{AoV}$ | 7.0 | $\text{g}/\text{cm}/\text{s}$ |
| tricuspid maximum opening angle | $\varphi_{TrV}^{\max}$ | $75^\circ$ | — |
| pulmonary maximum opening angle | $\varphi_{PuV}^{\max}$ | $75^\circ$ | — |
| mitral maximum opening angle | $\varphi_{MiV}^{\max}$ | $75^\circ$ | — |
| aortic maximum opening angle | $\varphi_{AoV}^{\max}$ | $75^\circ$ | — |
| tricuspid minimum opening angle | $\varphi_{TrV}^{\min}$ | (m)$5^\circ$ (s)$10^\circ$ | — |
| pulmonary minimum opening angle | $\varphi_{PuV}^{\min}$ | (m)$5^\circ$ (s)$10^\circ$ | — |
| mitral minimum opening angle | $\varphi_{MiV}^{\min}$ | (m)$5^\circ$ (s)$10^\circ$ | — |
| aortic minimum opening angle | $\varphi_{AoV}^{\min}$ | (m)$5^\circ$ (s)$10^\circ$ | — |

(m): mild AI condition.
(s): severe AI condition.

**Table 9:**
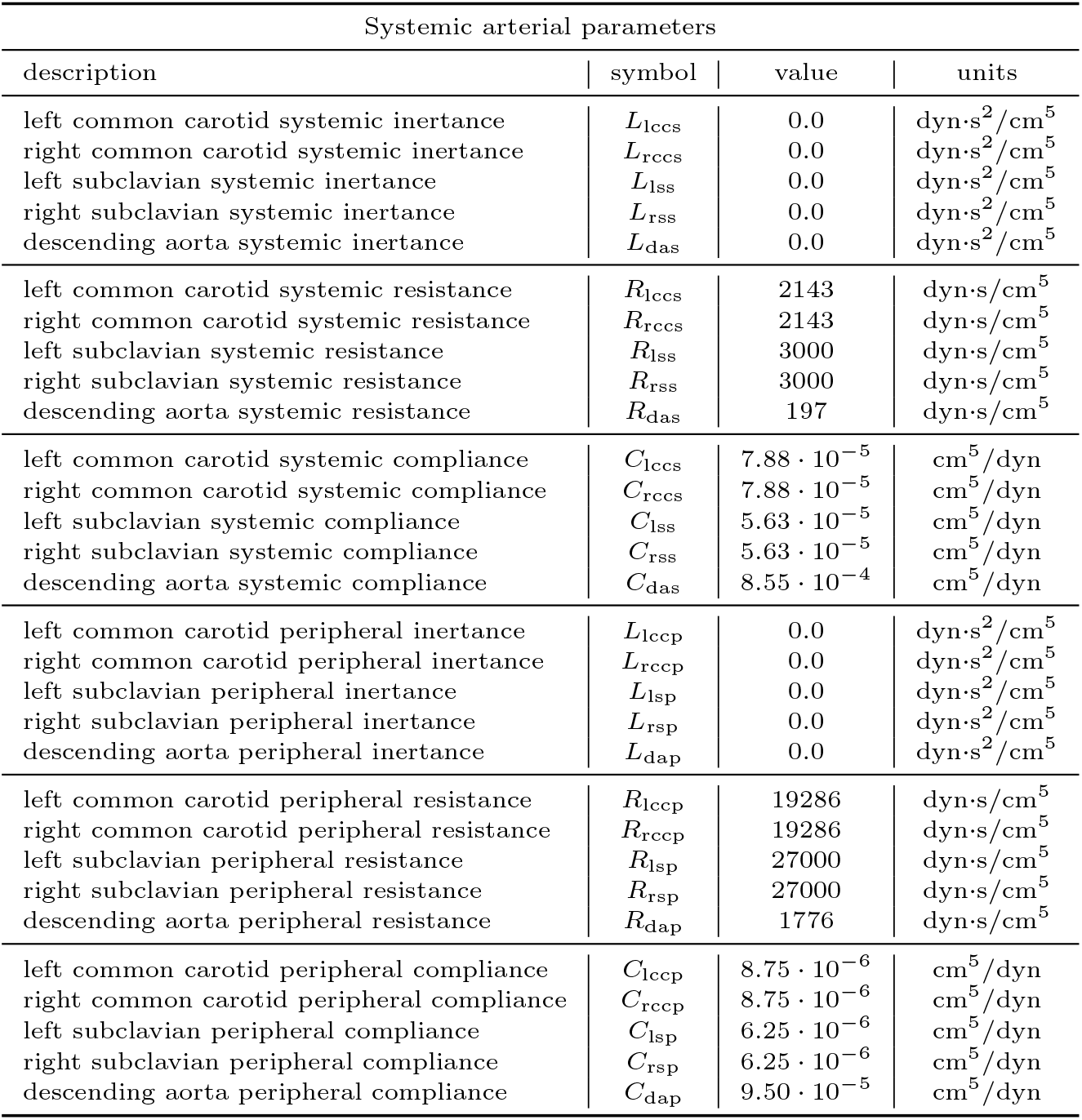
Description of model parameters that characterize the blood flow in the systemic arterial territories.

**Table 10:**
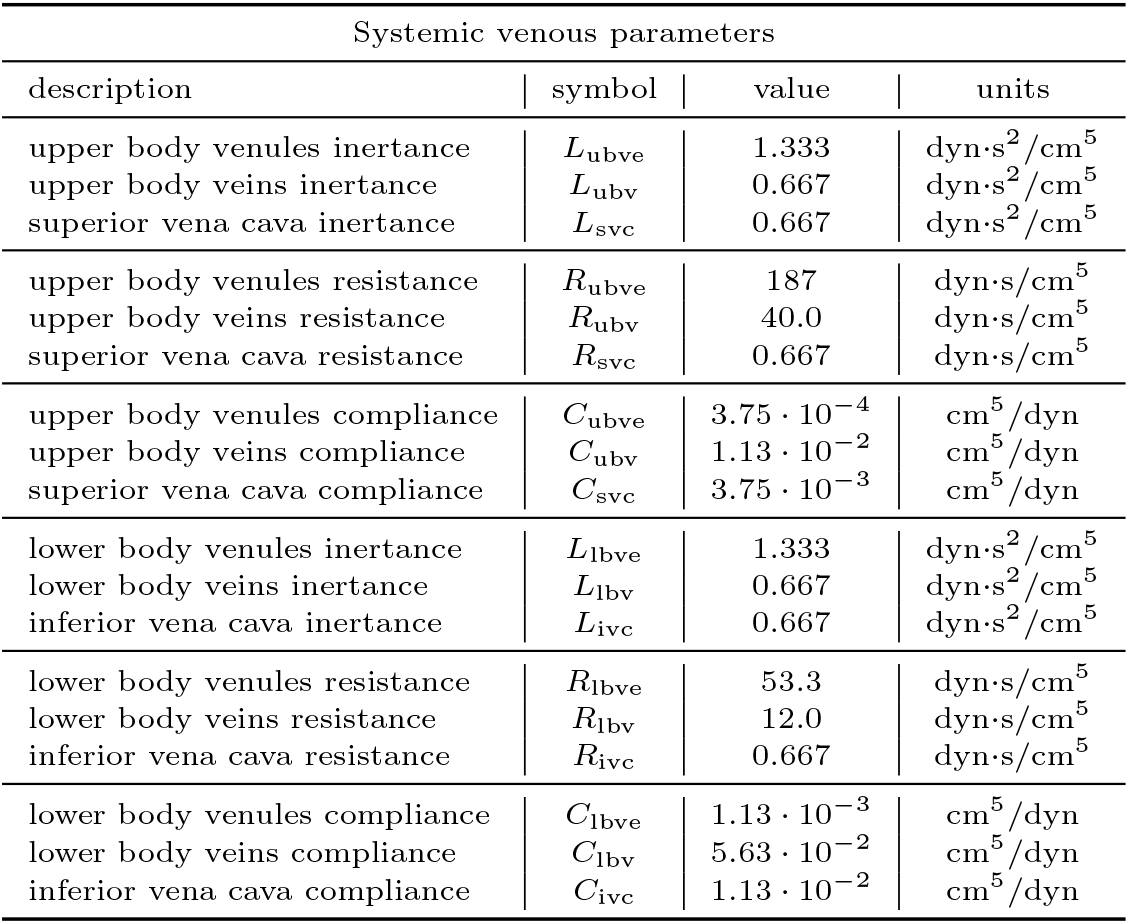
Description of model parameters that characterize the blood flow in the upper and lower venous regions.

**Table 11:** Description of model parameters that characterize the blood flow in the pulmonary arteries and veins

| Pulmonary parameters |  |  |  |
| --- | --- | --- | --- |
| description | symbol | value | units |
| pulmonary arteries inertance | $L_{pa}$ | 0.0 | $\text{dyn}\cdot\text{s}^2/\text{cm}^5$ |
| pulmonary veins inertance | $L_{pv}$ | 0.0 | $\text{dyn}\cdot\text{s}^2/\text{cm}^5$ |
| pulmonary arteries resistance | $R_{pa}$ | (c)160 (u)267 | $\text{dyn}\cdot\text{s}/\text{cm}^5$ |
| pulmonary veins resistance | $R_{pv}$ | (c)20 (u)33 | $\text{dyn}\cdot\text{s}/\text{cm}^5$ |
| pulmonary arteries compliance | $C_{pa}$ | (c)$2.32 \cdot 10^{-3}$ (u)$3.09 \cdot 10^{-4}$ | $\text{cm}^5/\text{dyn}$ |
| pulmonary veins compliance | $C_{pv}$ | (c)$2.25 \cdot 10^{-2}$ (u)$3.00 \cdot 10^{-3}$ | $\text{cm}^5/\text{dyn}$ |

(c): coupled RV condition.
(u): uncoupled RV condition.

**Table 12:** Description of additional model parameters that characterize closed-loop model coupled to the LVAD.

| LVAD parameters |  |  |  |
| --- | --- | --- | --- |
| description | symbol | value | units |
| LVAD speed | $S_{LVAD}$ | $\{5500, 6400\}^\dagger$ | rpm |

† : depending on the simulation protocol

The model parameters and boundary conditions for the local circulation model are described in Table 13.

**Table 13:**
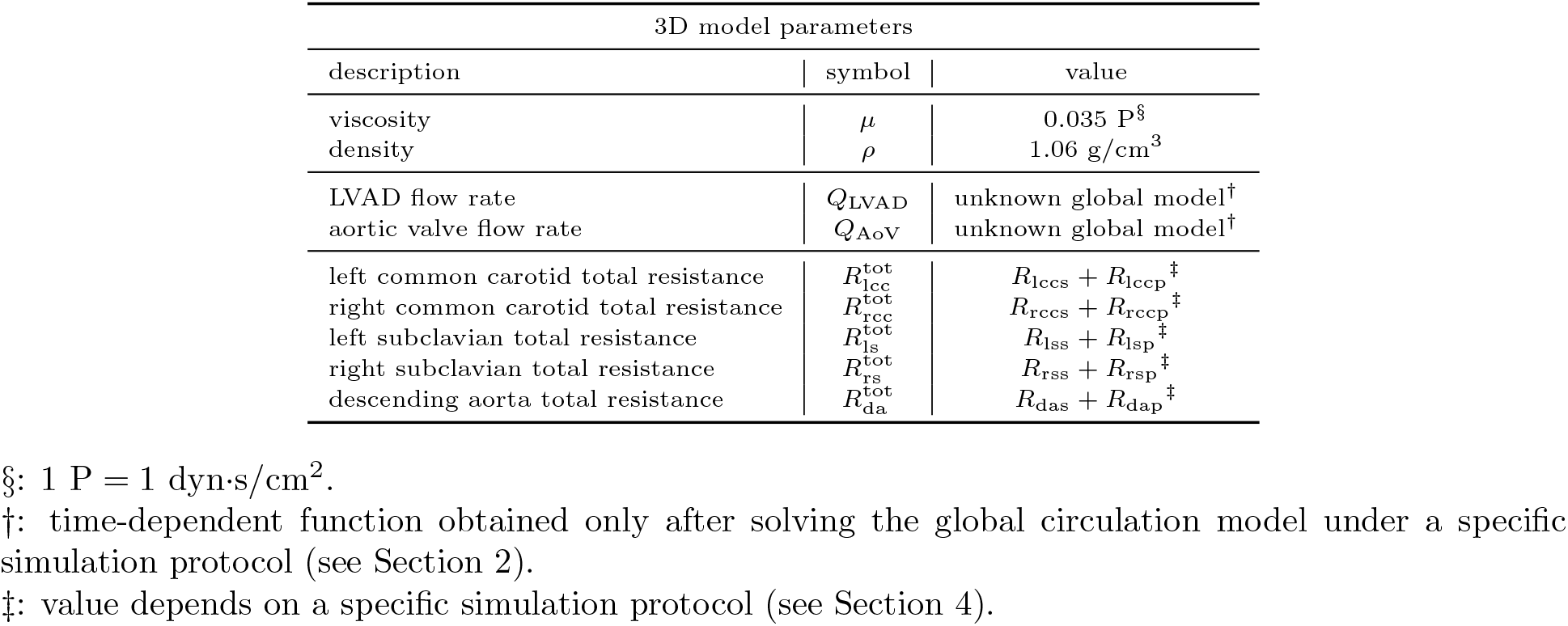
Description of model parameters that characterize the local circulation model.

### 4.2 .Cardiovascular conditions

Parameters reported in the tables described above are mostly shared by the two physiological conditions involving the right ventricle (RV) considered in the present study, these are: coupled and uncoupled RV. The RV was deemed to be uncoupled when the ratio of RV end-systolic elastance (*E*_es_) relative to the pulmonary effective arterial elastance (*E*_a_) was 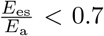 and coupled when the RV ratio satisfied 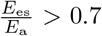. These two physiological conditions are differentiated in the tables by the preceding symbols (c) and (u), for coupled and uncoupled conditions, respectively.

Concerning aortic insufficiency (AI), we propose two different scenarios, these are: severe AI, which was defined as a regurgitant fraction (RF) of > 50%, and mild/moderate AI as a RF of *<* 50%. These two conditions are descrobed in the tables by the preceding symbols (s) and (m), for severe and mild AI, respectively.

For both, coupled and uncoupled RV conditions, we consider the following physiological scenarios:

- baseline (BL)
- blood pressure (BP) control
- speed augmentation (SA)
- pulmonary vasodilation (PV) for the uncoupled RV condition
- pulmonary vasodilation (PV-BP) and BP control for the uncoupled RV condition

The parameters that are modified in each scenario are described in Table 14.

**Table 14:** Description of multiplicative factors (except for the LVAD speed) that change the parameter values to simulate the physiological scenarios of interest.

| coupled RV – multiplicative factors (not for LVAD speed) |  |  |  |  |  |
| --- | --- | --- | --- | --- | --- |
| parameters | scenario |  |  |  |  |
|  | BL | BP | SA |  |  |
| $S_{LVAD}$ | 5500 rpm | 5500 rpm | 6400 rpm | | |
| $\{R_{lccs}, R_{rccs}, R_{lss}, R_{rss}, R_{das}\}$ | (s)2.0 (m)0.9 | (s)0.8 (m)0.5 | (s)1.5 (m)0.6 | | |
| $\{R_{lccp}, R_{rccp}, R_{lsp}, R_{rsp}, R_{dap}\}$ | (s)2.0 (m)0.9 | (s)0.8 (m)0.5 | (s)1.5 (m)0.6 | | |
| $\{C_{lccs}, C_{rccs}, C_{lss}, C_{rss}, C_{das}\}$ | 1.5 | 1.5 | 1.5 | | |
| $\{C_{lccp}, C_{rccp}, C_{lsp}, C_{rsp}, C_{dap}\}$ | 1.5 | 1.5 | 1.5 | | |
| $\{R_{pa}, R_{pv}\}$ | (s)2.0 (m)0.9 | (s)0.8 (m)0.5 | (s)1.5 (m)0.6 | | |
| $\{C_{pa}, C_{pv}\}$ | 1.5 | 1.5 | 1.5 | | |

| uncoupled RV – multiplicative factors (not for LVAD speed) |  |  |  |  |  |
| --- | --- | --- | --- | --- | --- |
| parameters | scenario |  |  |  |  |
|  | BL | BP | SA | PV | PV-BP |
| $S_{LVAD}$ | 5500 rpm | 5500 rpm | 6400 rpm | 5500 rpm | 5500 rpm |
| $\{R_{lccs}, R_{rccs}, R_{lss}, R_{rss}, R_{das}\}$ | (s)2.0 (m)0.9 | (s)1.5 (m)0.6 | (s)1.5 (m)0.6 | (s)2.0 (m)0.9 | (s)0.8 (m)0.5 |
| $\{R_{lccp}, R_{rccp}, R_{lsp}, R_{rsp}, R_{dap}\}$ | (s)2.0 (m)0.9 | (s)1.5 (m)0.6 | (s)1.5 (m)0.6 | (s)2.0 (m)0.9 | (s)0.8 (m)0.5 |
| $\{C_{lccs}, C_{rccs}, C_{lss}, C_{rss}, C_{das}\}$ | 0.8 | 1.3 | 0.8 | 1.5 | 1.5 |
| $\{C_{lccp}, C_{rccp}, C_{lsp}, C_{rsp}, C_{dap}\}$ | 0.8 | 1.3 | 0.8 | 1.5 | 1.5 |
| $\{R_{pa}, R_{pv}\}$ | (s)2.0 (m)0.9 | (s)1.5 (m)0.6 | (s)1.5 (m)0.6 | (s)1.2 (m)0.54 | (s)0.48 (m)0.3 |
| $\{C_{pa}, C_{pv}\}$ | 0.8 | 1.3 | 0.8 | 11.25 | 11.25 |

(m): mild AI condition.
(s): severe AI condition.

## 5. Results

Figure 5 shows the pressure-volume loops for both RV and LV in the case of a coupled RV. In that figure, the left column corresponds to the severe AI condition, and the right column to the moderate AI condition. The first row is the BL (baseline) scenario, the second row is the SA (speed augmentation) scenario and the third row is for the BP (blood pressure control) scenario. Analogously, Figure 6 shows the RV and LV pressure-volume loops in the case of an uncoupled RV. While the left column corresponds to the severe AI condition, the right column is for the moderate AI condition. The first, second and third rows are for the BL, SA and BP scenarios, respectively. Finally, fourth and fifth rows correspond to the PV (pulmonary vasodilation) and PV-BP (pulmonary vasodilation + blood pressure control) scenarios, respectively. Table 15 reports the indexes related to the cardiac function for both RV and LV in the case of a coupled RV. For the case of an uncoupled RV, these data are reported in Table 15.

**Table 15:** Indexes that characterize the cardiac function in the case of a coupled RV.

| Coupled RV |  |  |  |  |  |  |  |
| --- | --- | --- | --- | --- | --- | --- | --- |
| Cardiac index | Units | severe AI |  |  | moderate AI |  |  |
|  |  | BL | SA | BP | BL | SA | BP |
| $E_{esRV}$ | mmHg/ml | 0.364 | 0.360 | 0.356 | 0.364 | 0.351 | 0.354 |
| $E_{aRV}$ | mmHg/ml | 0.477 | 0.408 | 0.333 | 0.379 | 0.273 | 0.271 |
| $F_{RV}^{es/a}$ | - | 0.762 | 0.884 | 1.069 | 0.959 | 1.285 | 1.310 |
| $SW_{RV}$ | mmHg.ml | 851 | 860 | 827 | 862 | 805 | 816 |
| $MVO_{2RV}$ | mlO<sub>2</sub> | 0.0447 | 0.0440 | 0.0413 | 0.0435 | 0.0401 | 0.0398 |
| $PE_{RV}$ | mmHg.ml | 726 | 684 | 589 | 659 | 555 | 532 |
| $PVA_{RV}$ | mmHg.ml | 1577 | 1544 | 1416 | 1520 | 1360 | 1348 |
| $Eff_{RV}$ | - | 0.127 | 0.130 | 0.133 | 0.132 | 0.134 | 0.136 |
| $\bar{P}_{sa}$ | mmHg | 92 | 99 | 73 | 92 | 88 | 72 |
| $P_{sa}^{max}$ | mmHg | 97 | 101 | 78 | 94 | 89 | 75 |
| $P_{sa}^{min}$ | mmHg | 88 | 96 | 69 | 90 | 87 | 70 |
| $\bar{P}_{pa}$ | mmHg | 24 | 22 | 20 | 22 | 19 | 18 |
| $P_{pa}^{max}$ | mmHg | 28 | 28 | 26 | 27 | 25 | 25 |
| $P_{pa}^{min}$ | mmHg | 19 | 18 | 16 | 17 | 14 | 14 |
| $\Delta P_{AoV}$ | mmHg | 56.6 | 66.6 | 42.6 | 58.1 | 62.7 | 43.7 |
| $Q_{AoV}$ | l/min | -3.33 | -4.07 | -2.47 | -0.87 | -0.99 | -0.42 |
| $Q_{LVAD}$ | l/min | 5.42 | 7.05 | 6.26 | 5.35 | 7.20 | 6.22 |
| $V_{AoV}^{back}$ | ml | -59 | -68 | -50 | -15 | -16 | -12 |
| $Q_{AoV} + Q_{LVAD}$ | l/min | 2.10 | 2.97 | 3.80 | 4.47 | 6.21 | 5.80 |
| RF | - | 61.3% | 57.8% | 39.4% | 16.3% | 13.7% | 6.7% |

**Table 16:** Indexes that characterize the cardiac function in the case of an uncoupled RV.

| Uncoupled RV |  |  |  |  |  |  |  |  |  |  |  |
| --- | --- | --- | --- | --- | --- | --- | --- | --- | --- | --- | --- |
| Cardiac index | Units | severe AI |  |  |  |  | moderate AI |  |  |  |  |
|  |  | BL | SA | BP | PV | PV-BP | BL | SA | BP | PV | PV-BP |
| $E_{esRV}$ | mmHg/ml | 0.340 | 0.331 | 0.339 | 0.364 | 0.356 | 0.327 | 0.315 | 0.326 | 0.364 | 0.354 |
| $E_{aRV}$ | mmHg/ml | 1.192 | 0.979 | 0.918 | 0.477 | 0.333 | 0.857 | 0.585 | 0.597 | 0.379 | 0.271 |
| $F_{RV}^{es/a}$ | - | 0.285 | 0.338 | 0.370 | 0.762 | 1.069 | 0.381 | 0.540 | 0.547 | 0.959 | 1.310 |
| $SW_{RV}$ | mmHg.ml | 754 | 842 | 822 | 851 | 827 | 891 | 994 | 977 | 862 | 816 |
| $MVO_{2RV}$ | mlO<sub>2</sub> | 0.0602 | 0.0595 | 0.0578 | 0.0447 | 0.0413 | 0.0584 | 0.0530 | 0.0538 | 0.0435 | 0.0398 |
| $PE_{RV}$ | mmHg.ml | 1564 | 1445 | 1384 | 726 | 589 | 1345 | 988 | 1038 | 659 | 532 |
| $PVA_{RV}$ | mmHg.ml | 2318 | 2287 | 2206 | 1577 | 1416 | 2235 | 1982 | 2015 | 1520 | 1348 |
| $Eff_{RV}$ | - | 0.083 | 0.094 | 0.095 | 0.127 | 0.133 | 0.101 | 0.125 | 0.121 | 0.132 | 0.136 |
| $\bar{P}_{sa}$ | mmHg | 83 | 89 | 79 | 92 | 73 | 84 | 82 | 72 | 92 | 72 |
| $P_{sa}^{max}$ | mmHg | 88 | 92 | 82 | 97 | 78 | 86 | 82 | 73 | 94 | 75 |
| $P_{sa}^{min}$ | mmHg | 79 | 87 | 76 | 88 | 69 | 82 | 82 | 71 | 90 | 70 |
| $\bar{P}_{pa}$ | mmHg | 21 | 21 | 21 | 24 | 20 | 20 | 19 | 19 | 22 | 18 |
| $P_{pa}^{max}$ | mmHg | 41 | 39 | 39 | 28 | 26 | 38 | 34 | 34 | 27 | 25 |
| $P_{pa}^{min}$ | mmHg | 13 | 13 | 13 | 19 | 16 | 13 | 12 | 13 | 17 | 14 |
| $\Delta P_{AoV}$ | mmHg | 56.8 | 71.7 | 52.6 | 56.6 | 42.6 | 68.8 | 74.5 | 54.6 | 58.1 | 43.7 |
| $Q_{AoV}$ | l/min | -3.78 | -4.34 | -3.62 | -3.33 | -2.47 | -1.05 | -1.10 | -0.93 | -0.87 | -0.42 |
| $Q_{LVAD}$ | l/min | 5.56 | 6.92 | 5.80 | 5.42 | 6.26 | 5.05 | 6.86 | 5.80 | 5.35 | 6.22 |
| $V_{AoV}^{back}$ | ml | -63 | -72 | -60 | -59 | -50 | -17 | -18 | -15 | -15 | -12 |
| $Q_{AoV} + Q_{LVAD}$ | l/min | 1.78 | 2.59 | 2.19 | 2.10 | 3.80 | 4.00 | 5.77 | 4.87 | 4.47 | 5.80 |
| RF | - | 67.9% | 62.6% | 62.4% | 61.3% | 39.4% | 20.8% | 16.0% | 16.0% | 16.3% | 6.7% |

**Figure 5:**
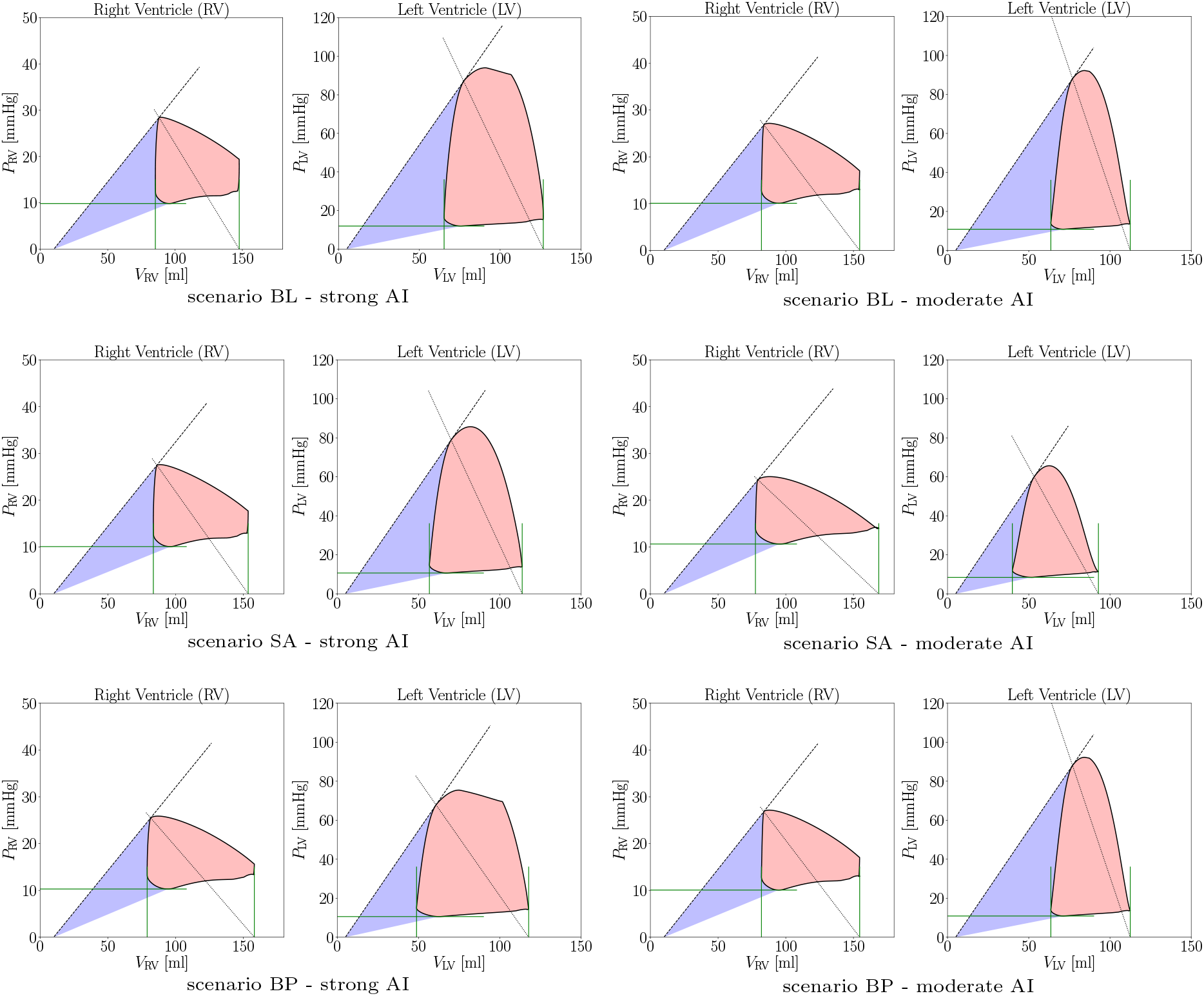
Pressure-volume loops in the case of a coupled RV for strong and moderate AI, and for the different physiological scenarios (BL, SA, BP).

**Figure 6:**
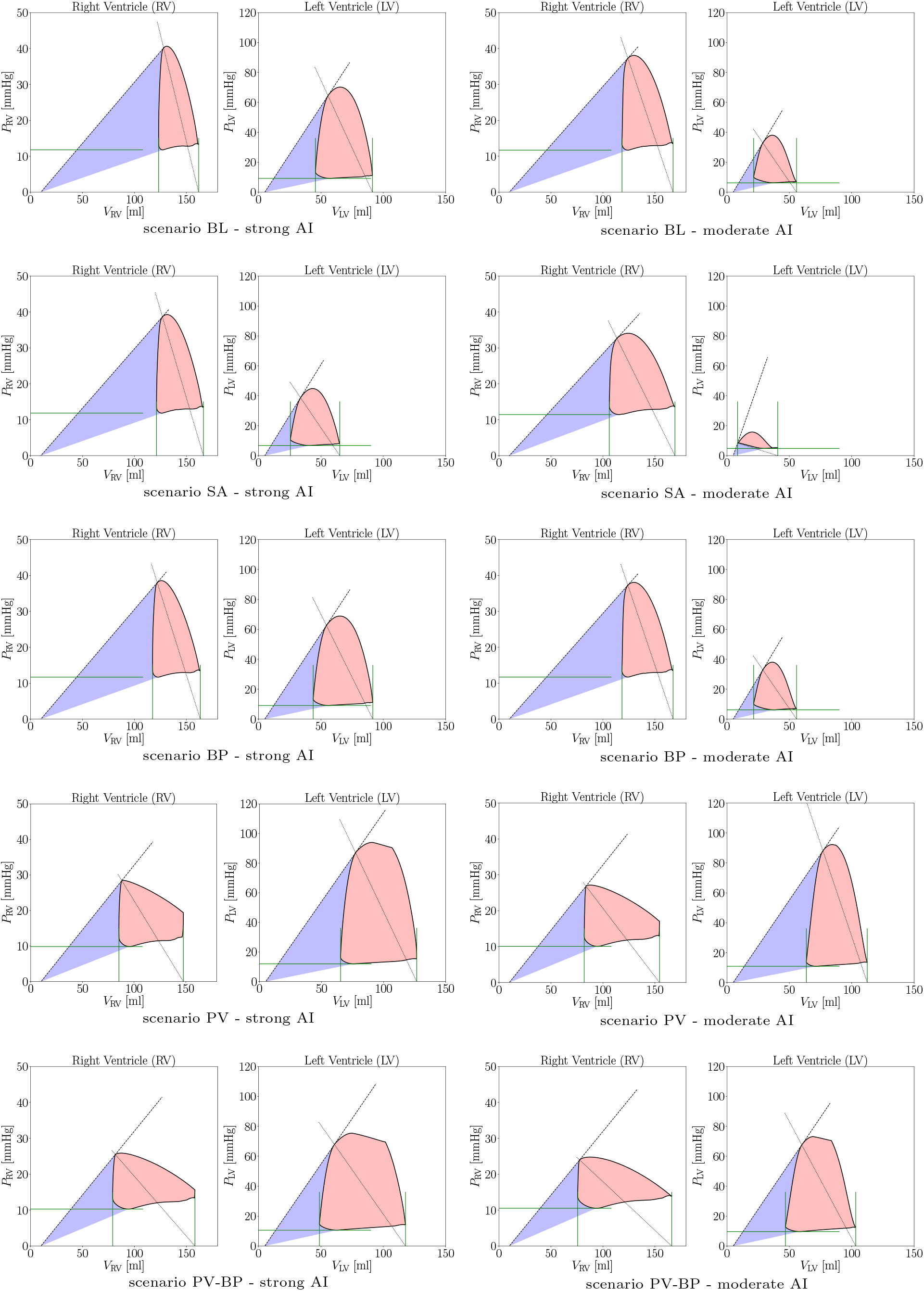
Pressure-volume loops in the case of an uncoupled RV for strong and moderate AI, and for the different physiological scenarios (BL, SA, BP, PV, PV-BP).

Figure 7 displays the evolution of some hemodynamic quantities along the cardiac cycle for the case of a coupled RV, and for strong and strong (top) and moderate (bottom) AI, respectively. In these figures, we report the RV and LV pressure and volume signatures, the systemic arterial and pulmonary arterial pressures, the left atrium and pulmonary veins pressure contours, the flow rate through the aortic root and through the LVAD, and the pressure-volume loops. Each figure features the hemodynamic signals for the three scenarios of interest, namely baseline (BL), speed augmentation (SA) and blood pressure (BP) control. Similar plots are reported in Figure 8 for the case of an uncoupled RV and strong (top) and moderate (bottom) AI, and for the five scenarios of interest, baseline (BL), speed augmentation (SA), blood pressure (BP) control, pulmonary vasodilation (PV) and pulmonary vasodilation plus blood pressure control (PV-BP).

**Figure 7:**
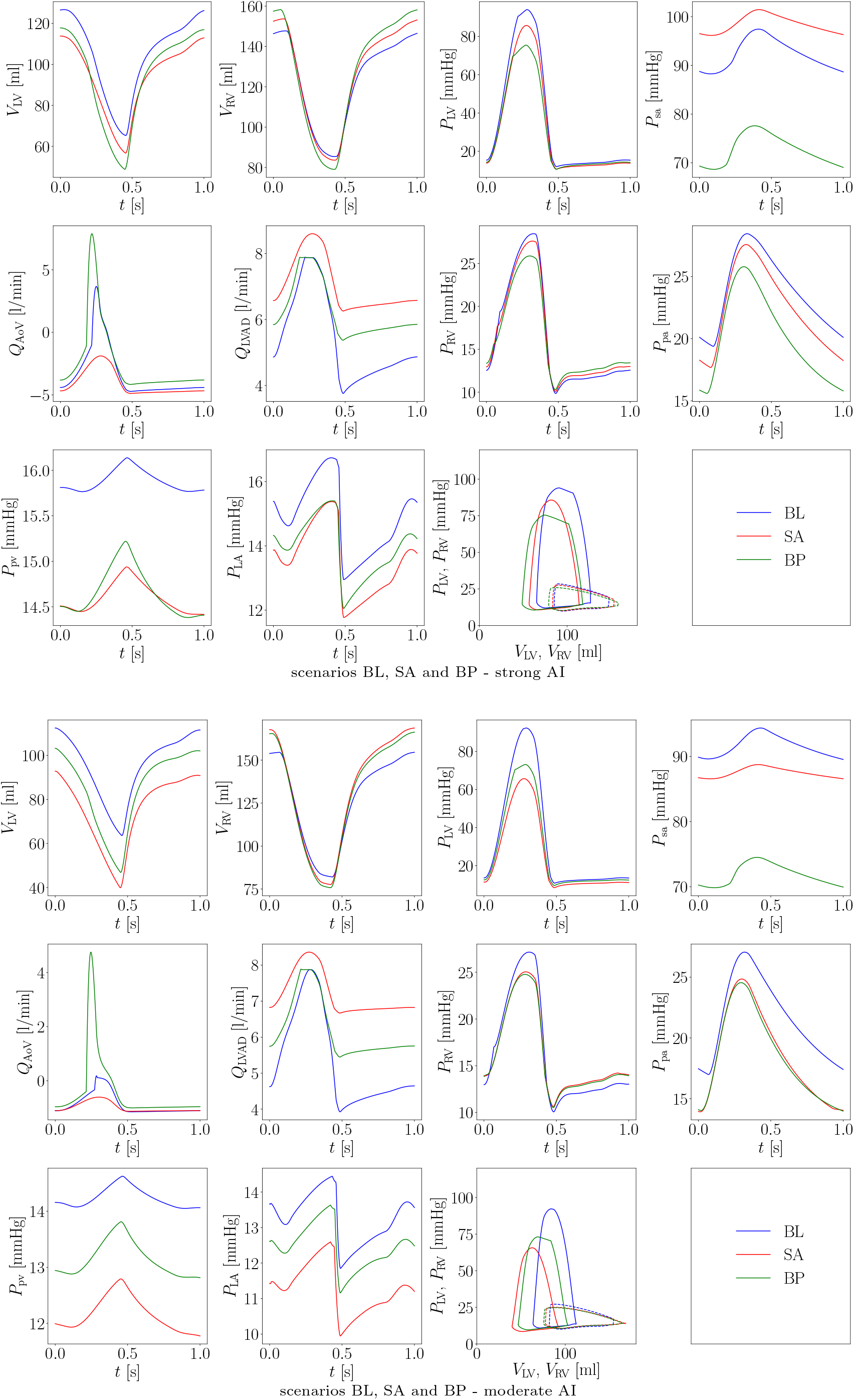
Pressure, volume and flow rate waveforms in the heart, systemic arteries and pulmonary vasculature for the case of a coupled RV. Top: condition with strong AI. Bottom: condition with moderate AI.

**Figure 8:**
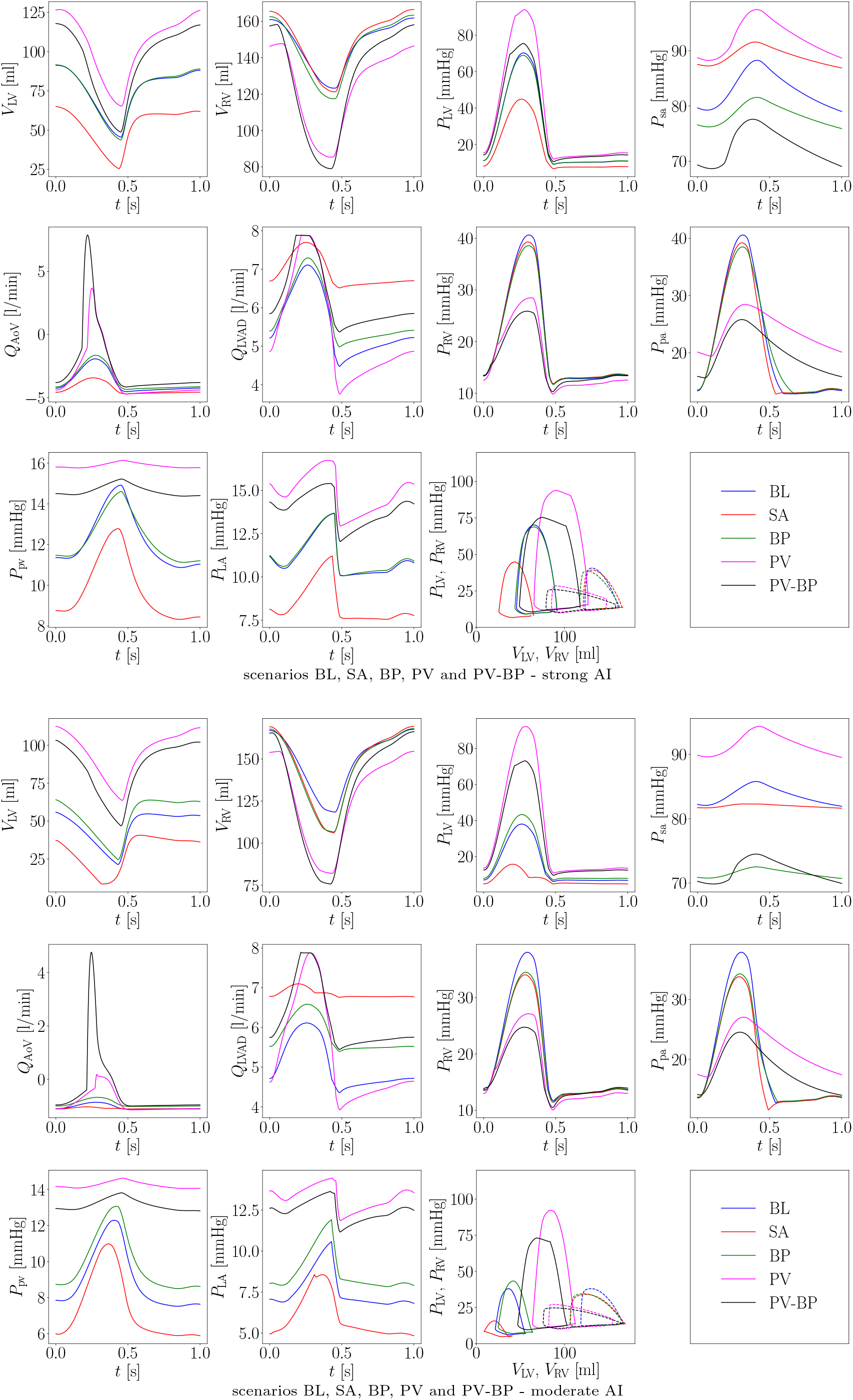
Pressure, volume and flow rate waveforms in the heart, systemic arteries and pulmonary vasculature for the case of an uncoupled RV. Top: condition with strong AI. Bottom: condition with moderate AI.

## References

[1] P. Blanco, R. Feijóo, A dimensionally-heterogeneous closed-loop model for the cardiovascular system and its applications, Medical Engineering and Physics 35 (5) (2013) 652–667.

[2] F. Liang, S. Takagi, R. Himeno, H. Liu, Multi-scale modeling of the human cardiovascular system with applications to aortic valvular and arterial stenoses, Medical and Biological Engineering and Computing 47 (7) (2009) 743–755.

[3] J. P. Mynard, Computer modelling and wave intensity analysis of perinatal cardiovascular function and dysfunction, Ph.D. thesis, The University of Melbourne (2011).

[4] T. Korakianitis, Y. Shi, Numerical simulation of cardiovascular dynamics with healthy and diseased heart valves, Journal of Biomechanics 39 (11) (2006) 1964–1982.

[5] T. Korakianitis, Y. Shi, A concentrated parameter model for the human cardiovascular system including heart valve dynamics and atrioventricular interaction, Medical Engineering and Physics 28 (7) (2006) 613–628.

[6] D. Burkhoff, K. Sagawa, Ventricular efficiency predicted by an analytical model, American Journal of Physiology - Regulatory Integrative and Comparative Physiology 250 (6 (19/6)) (1986) R1021–R1027.

[7] C. Bulant, P. Blanco, T. Lima, J. Assunção, A.N., G. Liberato, J. Parga, L. Ávila, A. Pereira, R. Feijóo, P. Lemos, A computational framework to characterize and compare the geometry of coronary networks, International Journal for Numerical Methods in Biomedical Engineering 33 (3).

[8] C. Bulant, P. Blanco, G. Maso Talou, C. Bezerra, P. Lemos, R. Feijóo, A head-to-head comparison between ct-and ivus-derived coronary blood flow models, Journal of Biomechanics 51 (2017) 65–76.

[9] A. Veneziani, C. Vergara, Flow rate defective boundary conditions in haemodynamics simulations, International Journal for Numerical Methods in Fluids 47 (8-9) (2005) 803–816.

